# Neural dynamics of visual streams interactions during memory-guided actions investigated by intracranial EEG

**DOI:** 10.1101/2024.08.20.608807

**Authors:** Sofiia Moraresku, Jiri Hammer, Vasileios Dimakopoulos, Michaela Kajsova, Radek Janca, Petr Jezdik, Adam Kalina, Petr Marusic, Kamil Vlcek

## Abstract

The dorsal and ventral visual streams play distinct roles in visual processing for action: the dorsal stream is assumed to support real-time actions, while the ventral stream facilitates memory-guided actions. As the recent evidence suggests a more integrated function of these streams, we investigated the neural dynamics and functional connectivity between them during memory-guided actions using intracranial EEG. We tracked neural activity in the inferior parietal lobule in the dorsal stream, and ventral temporal cortex in the ventral stream as well as hippocampus during a delayed action task. We found increased alpha power in both streams during the delay, indicating their role in maintaining visual information. We also observed an increase in theta band synchronization between the inferior parietal lobule and ventral temporal cortex, and between the inferior parietal lobule and hippocampus during the delay. Our study provides unique electrophysiological evidence for close interactions between dorsal and ventral streams, supporting an integrated processing model.

## Introduction

There has been an extensive debate on the precise role of the dorsal and ventral visual streams in visual processing for action. The original perception-action model, proposed by Goodale and Milner [1], implicates that the dorsal (in the occipito-parietal cortex) and ventral (in the occipito-temporal cortex) streams process visual information for different purposes: for motoric action and for conscious perception, respectively. The model also suggests that the two streams differ in storage capacity [2, 3]: the dorsal stream operates in real-time, storing visual information briefly for immediate action guidance, while the ventral stream appears to facilitate memory-guided or delayed actions. When visual information for action must be kept in memory for several seconds [2], the ventral stream becomes crucial for guiding the visuomotor act.

Evidence supporting the transient nature of the dorsal stream storage capacity is based on the studies in brain-damaged patients. For instance, patient D.F. with bilateral ventral lesions in the lateral occipital complex, demonstrated impaired object grasping after a short delay [3]. Conversely, patients with parietal lobe lesions and optic ataxia were less impaired after a short delay of 5 s compared to no delay [4], suggesting compensation by stored information in the ventral stream. Previous fMRI studies have also implicated that ventral stream regions play a role in delayed reaching and grasping [5], and transcranial magnetic stimulation inhibition of the lateral occipital complex in the ventral stream disrupted only delayed but not immediate grasping [6].

However, a recent review by Schenk and Hesse [7] argues that the ‘dorsal amnesia hypothesis’ oversimplifies the complex interactions between the dorsal and ventral streams in guiding actions. They discuss several fMRI studies showing activity in dorsal stream areas, such as the inferior parietal lobule (IPL), involved in maintaining target information for delayed reaching and grasping [8, 9]. It has been suggested that both streams might contribute to processing visual information for action in a more integrated manner. Results from one fMRI study seem to support this view, showing functional connectivity between occipitotemporal and parietal areas during sensorimotor tasks, suggesting a flexible, adaptive interaction between ventral and dorsal pathways [10]. However, no electrophysiological study has directly demonstrated this interaction during memory-guided actions. Moreover, previous studies have not fully addressed how different types of visual information, such as object identity and location, are integrated and utilized for delayed actions. The hippocampus may play an important role in this process as it appears to be involved in binding various features in episodic and working memory [11–13]. During memory-guided actions, the hippocampus may be involved in the integration of position and identity information together with the dorsal stream.

To address these gaps, we used intracranial EEG (iEEG) to investigate the dynamics of the dorsal and ventral streams during memory-guided actions based on different types of visual information. The high temporal resolution of iEEG allows to track neural activity simultaneously in the brain regions of the dorsal (IPL) and ventral (ventral temporal cortex, VTC) streams, as well as in the hippocampus, and study their interactions by means of functional connectivity. We focused on low-frequency theta and alpha activities due to their roles in working memory [14–17]. We analyzed functional connectivity using phase-locking value (PLV) between dorsal and ventral stream areas, as synchronized oscillations have been proposed as a mechanism for functional interactions between brain regions [18, 19]. These oscillations are considered to mediate long-range communication between cortical areas by low-frequency phase coupling [15, 20, 21].

Our study involved nine patients with drug-resistant epilepsy undergoing iEEG monitoring. They performed a delayed action task in a paradigm with two objects and a central cross. Their goal was to estimate which object was closer to the cross and to reach for it with a joystick after a jittered 4-second delay. We used two conditions: “same objects” and “different objects”, thus requiring attention to both object identity and location. Without any delay, this task is a simple egocentric distance estimation that primarily activates the parietal lobe in the dorsal stream (see review [22]). However, in the delayed task, we hypothesized the following: (1) sustained low-frequency activity in both the dorsal (IPL) and ventral (VTC) streams during the delay, based on the new framework of their combined role in processing visual information for memory-guided actions [7]; (2) increased synchronization between the IPL and VTC during the delay in both conditions, indicating integrated processing of spatial information; (3) additional activity in the hippocampus and stronger IPL-hippocampus synchronization during the delay in the “different objects” condition, which might integrate and maintain information about the object’s identity and position.

By addressing these hypotheses, our study aimed to advance the understanding of visual information processing in the brain, explore the neural basis of memory-guided actions, and further clarify the two visual streams model.

## Materials and Methods

### Patients

Nine patients (4 women; mean (± SEM) age = 37 ± 3 years, see details about subjects in Suppl. Table S1) with drug-resistant epilepsy were enrolled in our study at the Motol Epilepsy Center in Prague. They underwent iEEG monitoring for precise localization of the epileptic seizure onset zone before surgery. All the patients signed an informed consent to participate, and the study was approved by the Ethics Committee of Motol University Hospital. All the patients had normal or corrected to normal vision.

### Behavioral Task

The experimental task aimed to investigate the neural processes involved in memory-guided actions based on different types of visual information. Each trial consisted of fixation, encoding, delay, and action/recall phases. The trial began with a jittered fixation period of 1.9–2.1 s, displaying a white cross on a dark gray screen (Fig. 1A), followed by a 2 s encoding period with a central red cross and two types of objects: either two identical circles (“same” condition) or a square and a triangle (“different” condition). The positions of the objects were randomized for each trial, with one object always positioned closer to the cross than the other, maintaining a consistent distance ratio of 1.5. In the ‘different’ condition, the triangle was closer in 50% of the trials, and the square was closer in the other 50%. The participant’s task was to remember which object was closer to the cross: its position in the ‘same’ condition and both its position and identity in the ‘different’ condition. Since the cross was always centered on the screen, corresponding to the participant’s midsagittal plane, we assume that distance estimation was performed using egocentric coordinates.

**Fig. 1.**
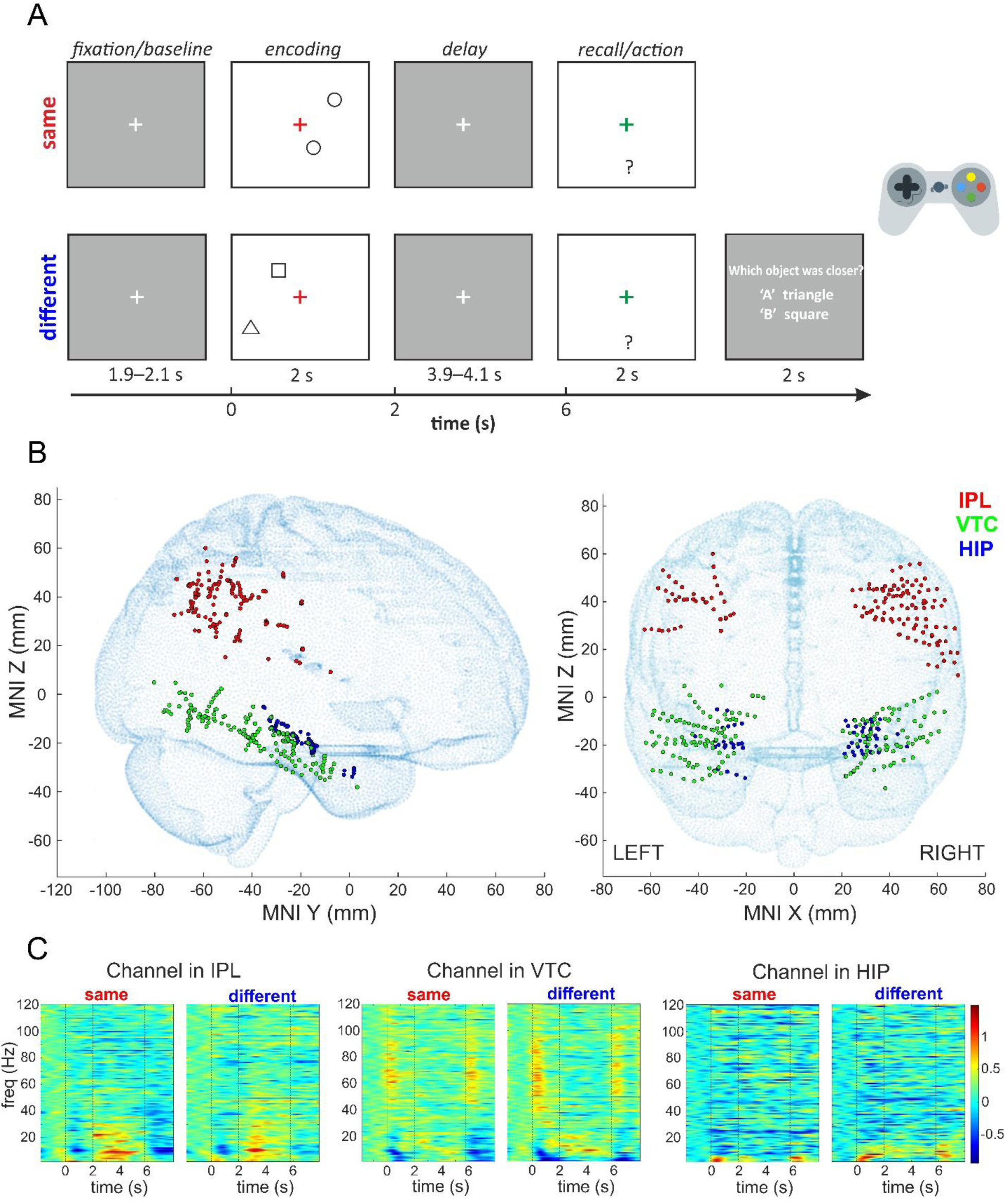
Task design, iEEG channel locations and exemplary responses. **A.** Experimental design of the delayed action task. Each trial included a jittered fixation period (1.9–2.1 s), encoding (2 s), a jittered delay (3.9–4.1 s), and an action/recall phase (2 s). Participants had to remember the object’s position (same condition) or both its position and identity (different condition) and reach for it with a joystick after the delay. In the different condition, two-alternative question about the object’s identity followed with 2 s time-window to answer. Participants responded using the gamepad: green ‘A’ for a triangle or red ‘B’ for a square. **B**. The projection of all implanted channels in 3 brain regions in 9 patients (369 channels) from both hemispheres on the standard MNI brain template (adult MNI-ICBM152 head model, [34]; http://www.ucl.ac.uk/dot-hub). The left and right panels show sagittal and coronal views, respectively. Legend: IPL, inferior parietal lobule; VTC, ventral temporal cortex; HIP, hippocampus. **C**. Trial-averaged spectrograms of exemplary individual channels in IPL, VTC, and HIP for the same and different conditions; 0 - the start of the encoding phase; vertical lines mark the boundaries of the task periods, and only the last 3.9 s of delay are plotted. Prominent changes during the delay can be observed in low-frequency power.

After the encoding phase, a white fixation cross reappeared on a dark gray screen during a delay phase, jittered between 3.9 and 4.1 s to prevent anticipatory responses. Following this, a green cross appeared for 2 s during the action/recall phase, prompting participants to respond. Using a gamepad joystick (Xbox wireless controller), they moved the green cross to the memorized position of the nearest object. The subjects made a straight reach from the center of the screen to the memorized object and returned to the center of the screen. A response was considered correct if the minimum distance of their trajectory was within a square area defined as the X,Y coordinates of the correct object plus or minus half the minimum distance between objects over all trials (10.5% of the screen width). Reaction time was measured from the start of the action phase to the time of reaching the correct area. In the ‘different’ condition, an additional two-alternative question about the object’s identity appeared for 2 s at the end of the trial. Participants responded by pressing a button on the gamepad: the green ‘A’ button for a triangle or the red ‘B’ button for a square. The total trial duration, including the pretrial period, ranged from 9.8 to 10.2 s for the ‘same’ condition and from 11.8 to 12.2 s for the ‘different’ condition.

The task was counterbalanced with 160 trials equally divided between the ‘same’ and ‘different’ conditions. Trials were grouped into blocks of 10, each assigned to one condition, with a subject-controlled break between blocks. The order of blocks was counterbalanced so that the average order of ‘same’ and ‘different’ blocks was the same. At the beginning of each block, participants received simple on-screen instructions about the upcoming condition. The experiment began with a brief presentation explaining the task, followed by a training session with shortened blocks of five trials per condition, with feedback provided after each trial. The experiment also included 160 immediate trials, in which participants reached for the object immediately without any delay, again equally divided between the ‘same’ and ‘different’ conditions. However, in this study, we only analyzed the delayed trials, focusing on the synchronization between two streams during the delay phase.

The experimental task was implemented in the PsychoPy3 v2020.1.3 [23] and run on a 15.6-inch TFT notebook monitor with a refresh rate of 60 Hz. The task and iEEG recording were synchronized by TTL pulses sent to a trigger port of an iEEG recording system at the start of each trial.

### Intracranial EEG Recording and Preprocessing

The iEEG was recorded with stereotactically implanted multi-contact electrodes, often also referred to as stereo-EEG. Recording sites were selected on an individual basis, strictly according to the medical requirements of the pre-surgical evaluation of epileptic zones, without reference to the present study. Eleven to fifteen semi-rigid electrodes were implanted intracerebrally per patient, depending on the suspected origin of their seizures. Each electrode had a diameter of 0.8 mm and consisted of eight to 18 contacts, 2 mm long and 1.5 mm apart (DIXI Medical Instruments). The iEEG signal was recorded with medical amplifiers (Quantum, NeuroWorks), sampled at 2048 Hz, and later downsampled to 512 Hz to reduce the computational load. The reference contact for each patient was located in the white matter. Postimplantation CT, co-registered with preimplantation MRI, was used to identify the positions of electrode contacts in each patient [24]. The anatomical positions of the electrode contacts were labeled by an experienced neurologist. The contact positions were then normalized to the Montreal Neurological Institute (MNI) space using standard Statistical Parametric Mapping algorithms (SPM 12) for group-level visualizations.

The downsampled iEEG recordings were first visually inspected, and any bad electrode contacts with obvious artifacts were discarded. Contacts identified as being in the seizure onset zone or heterotopic cortex were also excluded. Bipolar derivations were calculated between adjacent contacts to suppress contributions from distant neuronal assemblies; bipolar iEEG signals were visualized at the center between two contacts. Henceforth, we will refer to the bipolar contact pair simply as a ‘channel’. When the channel was derived from two contacts in a different brain structure, we labeled it with the structure with a larger unilateral response. Line noise was removed from the iEEG signal with a notch filter (4rd order Butterworth stop-band filter of 1 Hz width centered at the 50 Hz and harmonics, zero phase shift). Data processing and analysis were performed in Matlab R2018a.

For further analysis, we selected channels located in three brain regions of interest (containing a total of 369 channels in 9 patients, see Table 1 and Fig. 1B): IPL, including supramarginal and angular gyri and the adjacent lateral wall of intraparietal sulcus; VTC, including inferior temporal, lingual, fusiform and parahippocampal gyri; and the hippocampus (HIP).

**Table 1.**
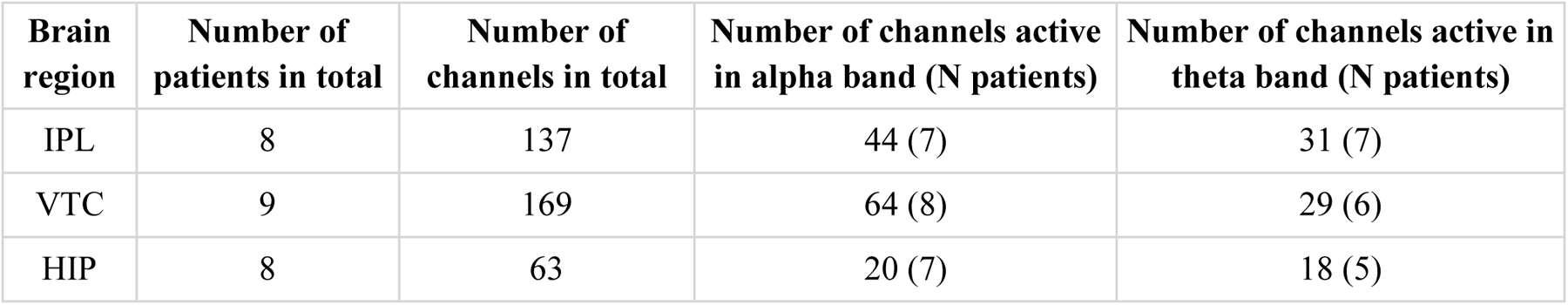
Number of implanted and active channels in each brain region.

| **Brain region** | **Number of patients in total** | **Number of channels in total** | **Number of channels active in alpha band (N patients)** | **Number of channels active in theta band (N patients)** |
| --- | --- | --- | --- | --- |
| IPL | 8 | 137 | 44 (7) | 31 (7) |
| VTC | 9 | 169 | 64 (8) | 29 (6) |
| HIP | 8 | 63 | 20 (7) | 18 (5) |

### Time-frequency analysis

Time-frequency analysis was performed for linearly increasing frequencies between 2 and 120 Hz, with a resolution of 1 Hz frequency bins, using a technique called filter-Hilbert method. Similar to our previous studies [25–27], we estimated the power change using the following procedure. First, we band-pass filtered the entire recording dataset (third-order Butterworth filter, zero phase shift) in consecutive non-overlapping 1 Hz frequency bands. For each band, we extracted the amplitude envelope using the Hilbert transform; the obtained envelope was downsampled to 64 Hz, resulting in a time resolution of 15.625 ms. The envelope of each band was normalized by dividing it by its mean value over the entire recording session, channel-wise for each frequency band, effectively whitening the broad frequency band and compensating for the 1/f frequency decay of EEG signals [28]. Then, similar to [16] and based on visual inspection of time-frequency spectrograms, we extracted power for two non-overlapping frequency bands: theta (2-7 Hz) and alpha (8-13 Hz). To do this, the original 1 Hz bands in these theta and alpha ranges were averaged and multiplied by 100 to obtain a single time series of theta and alpha power for each channel expressed in the percentages of the mean value. These two time series signals were then divided into epochs. To ensure that the epochs had the same duration, we left out from the epochs the jittered period from the beginning of the delay phase (0.0-0.2s). Each epoch was then divided into the following five task periods: (1) baseline (0.5 s, end of the fixation period), (2) encoding (2.0 s), (3) first half of delay (1.9 s, referred to as Delay 1 in the text), (4) second half of delay (1.9 s, referred to as Delay 2 in the text) and (5) recall (2.0 s). In the result, the epochs were from -500 to 7800 ms relative to the stimulus onset - the beginning of the encoding phase.

The mean of the -500 to 0 ms of the fixation period (i.e. the baseline) was subtracted from each epoch to remove signal changes independent of the respective stimulus. We excluded epochs in each channel containing interictal epileptiform discharges, which were identified by a spike detector implemented in MATLAB from further analysis [29]. Additionally, trials with incorrect behavioral responses—incorrect joystick responses for the object’s position and incorrect button responses for the object’s identity—were excluded from the iEEG analysis.

To test for the significance of the mean power in a given frequency band relative to the baseline, we used a Wilcoxon signed-rank test, with FDR correction across the time samples and channels, similar to our previous studies [25–27]. For each channel, we compared the average power for all trials of the respective condition during the pre-stimulus interval (-500– 0 ms before encoding) with all the time points during the post-stimulus period (0–7800 ms). As a conservative estimate, we used a sliding window of six samples (93.75 ms) with the highest p-value. If there was a significant difference at any time point relative to the baseline for a selected condition, the channel was considered **’active’** in the given frequency band.

Then, for all active channels in a given frequency band (theta or alpha), we averaged the power for each task period: encoding, first and second halves of delay, and recall. These averaged values were submitted to a linear mixed effects model (LMEM) with task period and condition and their interaction as fixed effects and channel and patient as random effects (p < 0.008, Bonferroni correction for six models - three brain regions and two frequency bands):

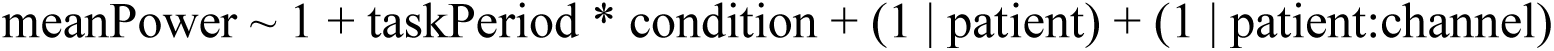

### Phase-locking value analysis

To evaluate the functional connectivity between channel pairs, we calculated the phase-locking value (PLV) [30]. This was done using a multitaper frequency transformation with two tapers based on the Fourier transform, covering a frequency range of 2-20 Hz with a resolution of 1 Hz, as implemented in the FieldTrip toolbox [31], and similar to other studies [15, 32].

The PLV between channels i and j is defined as:

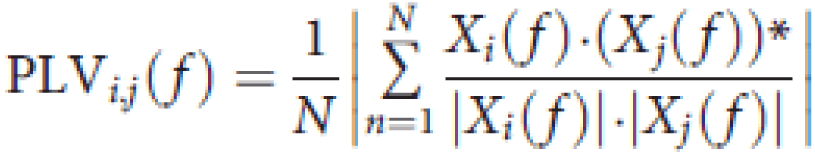

where N is the number of trials, X(f) is the Fourier transform of signal x(t) (1.9 s task period), and (·)* represents the complex conjugate.

Phase differences were calculated for each channel pair (i,j) between IPL and VTC, and between IPL and HIP, using the spectra of the two halves of the delay and recall periods to quantify inter-electrode phase coupling. To determine the statistical significance of PLV during both parts of the delay and recall, we compared them to the baseline (1.9-s fixation interval) using permutation statistics with cluster correction. The null distribution was created by calculating the PLV differences between the baseline and each task period (the first half of delay (1.9 s, Delay 1), the second half of delay (1.9 s, Delay 2), and recall (0.5 s, a part before the response was made). Trials were randomly assigned to these two periods, and the PLV differences between these periods were calculated and repeated 200 times. Only those frequency bins with PLV above the 95th percentile threshold were considered statistically significant [15, 32]. Cluster correction was applied to account for multiple comparisons across the 2-20 Hz frequency range (p < 0.05 was used to obtain null-hypothesis clusters). A channel pair was considered significant if its PLV was significant at any frequency point in the 2-20 Hz range after cluster correction.

We found that different channel pairs exhibited PLV significance at various frequency points within the 2-20 Hz range. We aggregated all channel pairs significant in any of the three task periods (two parts of the delay and recall) from all patients and calculated the ratio of significant pairs (Sig.P.Ratio) in each 1 Hz frequency bin. This was done by dividing the number of significant pairs between two brain regions by the total number of channel pairs between these regions, for all subjects and in each 1 Hz frequency bin independently [33]. This Sig.P.Ratio was calculated separately for IPL-VTC and IPL-HIP pairs. To identify frequency ranges at which the proportion of channel pairs was significantly above the chance level, we applied the binomial test to these Sig.P.Ratio values (one-sided test, p < 0.05, FDR-corrected across frequency bins). The median Sig.P.Ratio of each region pair across all frequency bins was used as the chance level for the binomial test [33].

We then averaged the PLV for each task period (baseline, two parts of the delay, and recall) for each frequency range with a Sig.P.Ratio significantly higher than the chance median level. These averaged values were then analyzed using the LMEM with task period, condition and their interaction as fixed effects, and channel pair and patient as random effects (p < 0.0167, Bonferroni correction for three models, each for one identified frequency range, see Results):

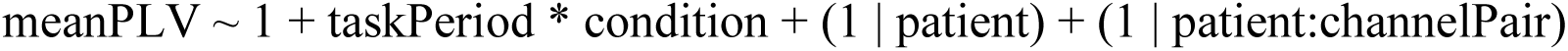

## Results

### Task Performance

All 9 patients completed the task (80 trials for the same condition and 80 trials for the different condition; the training session was not included in the analysis). There was no significant difference in reaction time and accuracy of joystick responses between the same and different conditions (Wilcoxon signed-rank test, p=0.4 for reaction time and p=0.53 for accuracy). The mean ± SEM reaction time and accuracy for the same condition were 813 ± 47 ms and 98.1 ± 0.7%, respectively. For the different condition, the mean ± SEM reaction time and accuracy were 801 ± 49 ms and 97.6 ± 1%, and the mean ± SEM accuracy for the object’s identity question was 91 ± 4.3%.

### Increased alpha activity during the delay in the dorsal and ventral streams

Based on visual analysis of time-frequency spectrograms (Fig. 1C), prominent changes during the delay were found in the alpha band. Mean alpha power 8-13 Hz was extracted for the whole length of the epoch (-500-7800 ms) for the same and different conditions and compared with the baseline (500 ms of pre-trial fixation period) to find active channels in this frequency band (see Methods section for details, an exemplary channel shown in Fig. 2A). Then, the mean alpha power across each task period and both conditions from all active channels in three brain regions (a total of 128 channels, see Table 1) from 9 patients was tested for significance by the LMEM (one model for each brain region).

**Fig. 2.**
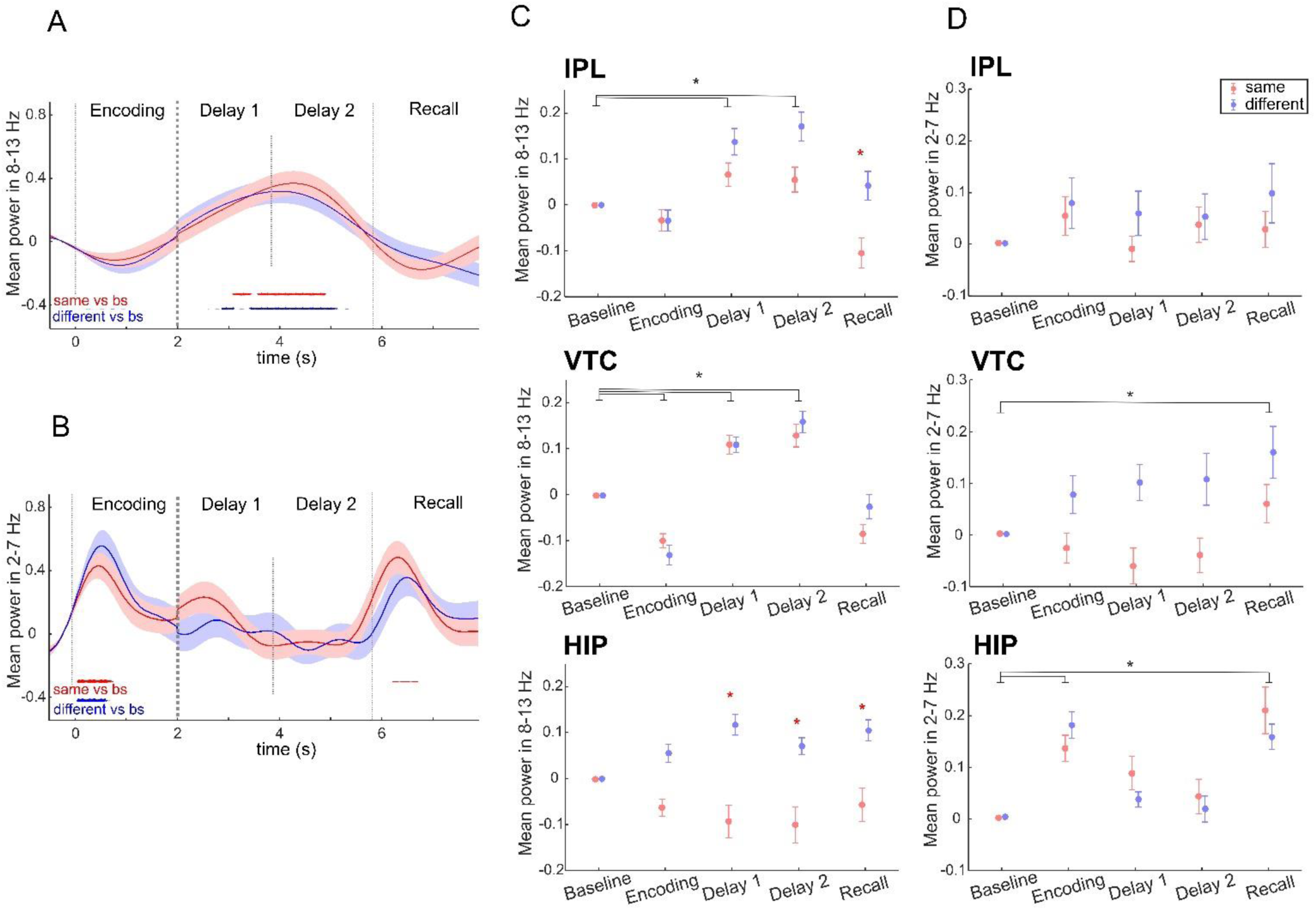
Time-frequency analysis. **A**. Example of an active channel in the VTC in the alpha band (8-13 Hz). The graph shows the mean ± SEM of power as a percentage of baseline activity. Mean responses to the “same” condition are shown in red, while mean responses to the “different” condition are shown in blue. Vertical lines mark the boundaries of the task periods; the last 3.9 s of delay are shown, without the 0.0-0.2s jitter at the beginning. Significant time points as compared to the baseline period are marked by asterisks (red for the same condition, blue for the different condition) using the FDR-corrected Wilcoxon rank sum test at p < 0.05. **B.** Example of the active channel in the HIP in the theta band (2-7 Hz). Same conventions as in panel A. **C.** Mean ± SEM across channels of alpha power (8-13 Hz) for each task period in three brain regions: IPL (44 channels), VTC (64 channels), and HIP (20 channels). The “same” condition is shown in red, and the “different” condition is shown in blue. The black asterisk indicates significant differences between task periods, and the red asterisk indicates significant differences between conditions within the same task period (p < 0.008, LMEM). **D.** Mean ± SEM across channels of theta power (2-7 Hz) for each task period in three brain regions: IPL (31 channels), VTC (29 channels), and HIP (18 channels). Same conventions as in panel C. Alpha power increased during the delay in both dorsal and ventral streams, while theta power increased only in the ventral stream during the recall and encoding.

The LMEM showed (Fig. 2C, see Suppl. Table S2 for further details) that in the IPL and VTC, alpha power increased for both conditions during the two parts of the delay compared to the baseline (*IPL*: main effect of task period Delay 1 t(430) = 4.16, p < 0.001, task period Delay 2 t(430) = 5.18, p < 0.001; *VTC*: task period Delay 1 t(630) = 4.46, p < 0.001, task period Delay 2 t(630) = 6.48, p < 0.001). In the IPL, there was also a difference between the same and different conditions, with the same condition showing a decrease in alpha power during recall (interaction between task period Recall and condition t(430) = 3.14, p = 0.002). In the HIP, alpha power increased only for the different condition during the two parts of the delay and recall (interaction between task period Delay 1 and condition t(190) = -4.57, p < 0.001, interaction between task period Delay 2 and condition t(190) = -3.73, p < 0.001, interaction between task period Recall and condition t(190) = -3.51, p < 0.001)).

### Increased theta power in the ventral stream

We also extracted mean theta power 2-7 Hz for the entire length of the epoch. Similar to the alpha power analysis, we identified active channels in this frequency band (an exemplary channel shown in Fig. 2B). We averaged the 2-7 Hz power across each task period and each condition from all active channels in three brain regions (a total of 78 channels, see Table 1) from 9 patients and tested the effect of task period and condition for significance using the LMEM.

The LMEM showed (Fig. 2D, see Suppl. Table S3 for further details) that, in the IPL, there was no significant difference in 2-7 Hz power between conditions and task periods or their interactions. In the VTC, theta power increased for both conditions during recall compared to the baseline (main effect of task period Recall t(280) = 3.51, p < 0.001). In the HIP, theta power also increased for both conditions during the recall, but also during the encoding (main effect of task period Recall t(170) = 4.91, p < 0.001, main effect of task period Encoding t(170) = 5.63, p < 0.001). In the VTC and HIP, the main effect of condition and the interaction between each task period and condition were not significant.

### Functional coupling between the dorsal and ventral streams

To investigate whether the dorsal and ventral streams communicate during the delay and recall phases in memory-guided actions, we first calculated the PLV across all trials for each channel pair between the IPL and VTC, and between the IPL and HIP within the same hemisphere for the first and second halves of the delay and for the recall phase. The number of channels and patients in each brain region included in the PLV analysis is shown in Table 2.

**Table 2.** Information about patients and channels used in the PLV analysis.

| Patient | IPL - VTC,<br>Hemisphere | Channels<br>(IPL/VTC) | IPL - HIP,<br>Hemisphere | Channels<br>(IPL/HIP) |
| --- | --- | --- | --- | --- |
| P1 | R | 19/18 | R | 19/5 |
| P2 | - | - | - | - |
| P3 | R | 13/19 | R | 13/3 |
| P4 | - | - | R | 4/11 |
| P5 | L | 6/34 | R | 7/9 |
| P6 | L | 7/15 | L | 7/8 |
| P7 | R | 33/4 | - | - |
| P8 | R | 6/16 | - | - |
| P9 | L | 21/44 | L | 21/5 |

Next, we identified the frequency ranges with most prominent connectivity for each pair of brain regions. First, to determine the statistical significance of PLV at each frequency bin, the PLV spectra for each period were compared to the baseline (see Methods section for details). We found that different channel pairs exhibited significant PLV at various frequency points within the 2-20 Hz range (Fig. 3). Therefore, in the second step, we identified the frequencies with the proportions of significant channel pairs for each brain region pair above the chance level. To this end, we compared the ratios of significant pairs (Sig.P.Ratios) between the regions at each frequency point with the median Sig.P.Ratio across all frequency points in the region pair using binomial tests (see Methods section for details). For IPL-VTC connections, the above-chance proportions of significant channel pairs were in the 2-5 Hz range (p < 0.05, FDR-corrected, Fig. 4A). For IPL-HIP connections, the above-chance proportions of significant channel pairs were identified in the 2-4 Hz and 7-8 Hz ranges (Fig. 4B).

**Fig. 3.**
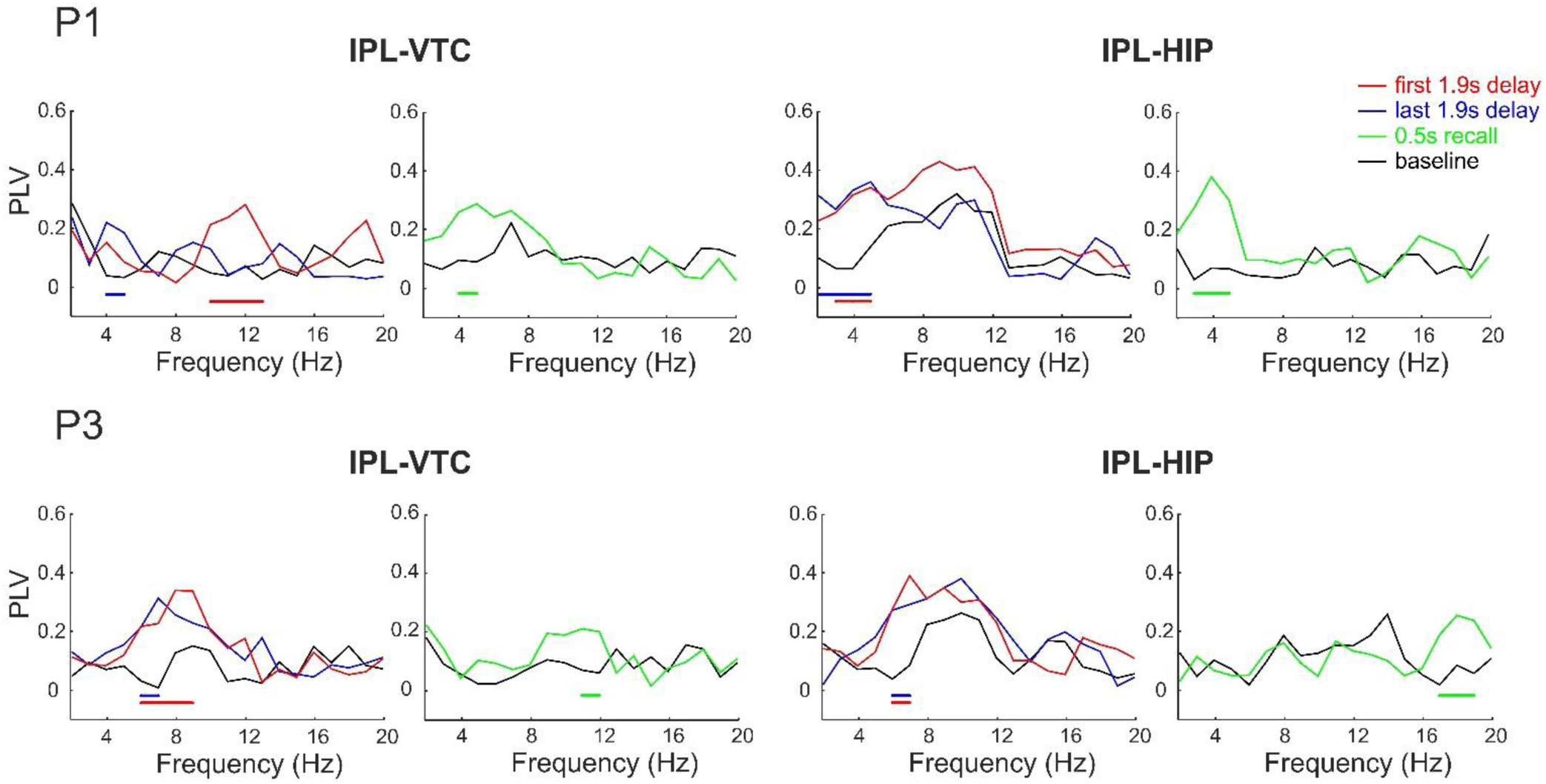
PLV spectra for individual channel pairs between IPL and VTC, and between IPL and HIP during baseline (black), first half of the delay (red), second half of the delay (blue), and 0.5 s of the recall phase (green) (examples for patients P1 and P3). Horizontal colored bars indicate frequency bins of significant PLV differences from baseline (p < 0.05, cluster-based non-parametric permutation test): red for the first half of the delay, blue for the second half, and green for the recall. Different channel pairs exhibited significant PLV differences from baseline at various frequencies within the 2-20 Hz range.

**Fig. 4.**
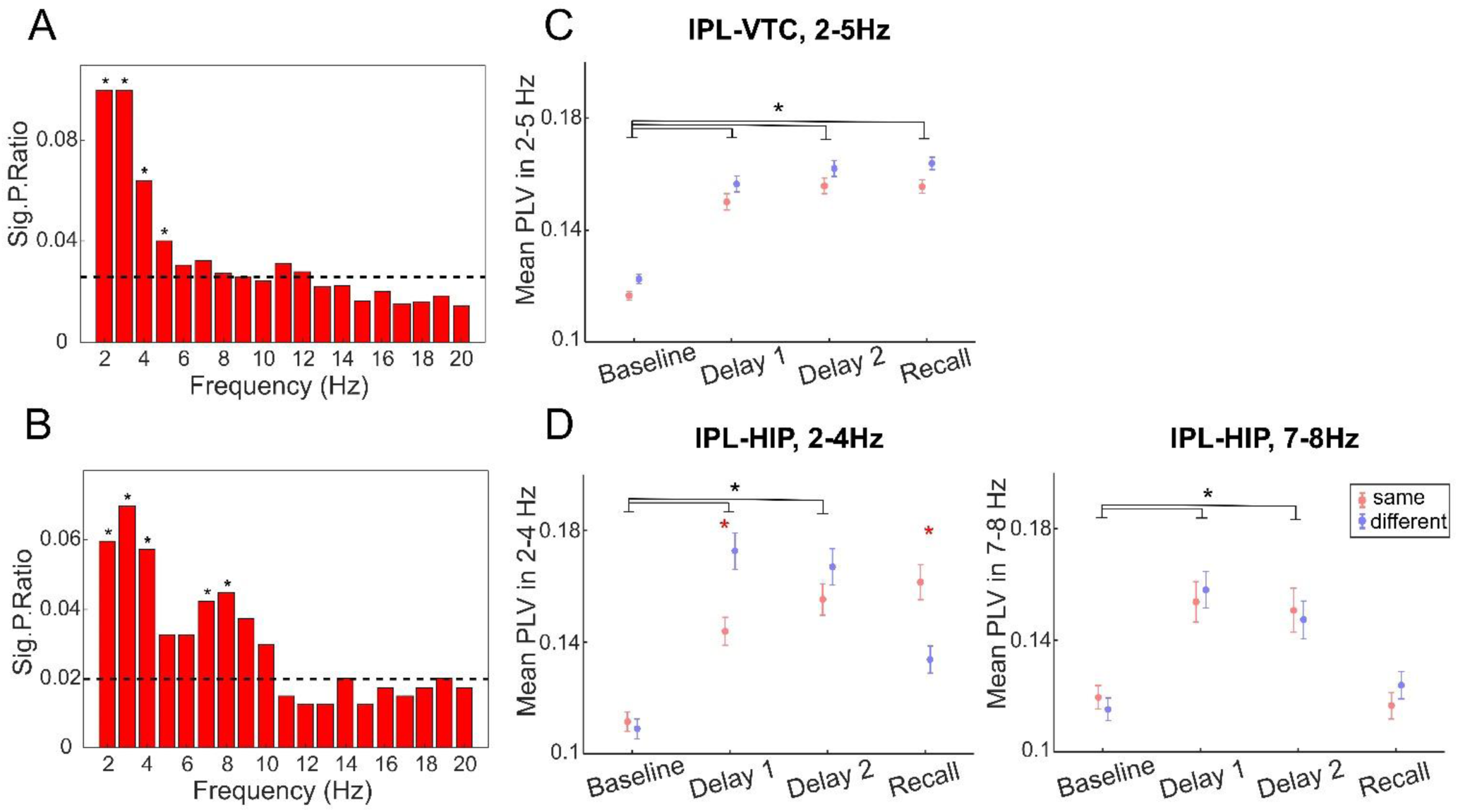
PLV analysis. **A.** Sig.P.Ratio between IPL and VTC across frequency bins (1 Hz intervals). The Sig.P.Ratio represents the proportion of channel pairs across all patients exhibiting significant PLV in each frequency bin. The dashed black line indicates the median Sig.P.Ratio across all frequency bins. Asterisks mark frequency bins with significantly higher Sig.P.Ratio compared to the median (binomial test, p < 0.05). **B.** Sig.P.Ratio between IPL and HIP across frequency bins (1 Hz intervals). The legend and description are consistent with panel A. **C.** Means and SEMs of PLV in the 2-5 Hz range for each task period between IPL and VTC (755 channel pairs). The “same” condition is depicted in red and the “different” condition in blue. The black asterisks denote significant differences between task periods, while the red asterisks mark significant differences between conditions within the same task period (p < 0.0167, LMEM). **D.** Means and SEMs of PLV in the 2-4 Hz and 7-8 Hz ranges for each task period between IPL and HIP (134 channel pairs). The legend and description are consistent with panel C. Significant increases in PLV were observed between IPL and VTC in the 2-5 Hz range and between IPL and HIP in the 2-4 Hz and 7-8 Hz ranges during delay, suggesting functional coupling between these regions. Notably, IPL-HIP connections showed condition-specific differences in PLV in the 2-4 Hz range during the first half of the delay and recall.

### Functional coupling between IPL and VTC

For all significant IPL-VTC connections (755 channel pairs in 7 patients), we calculated the PLV separately for each condition and averaged the PLV over the 2-5 Hz range for each task period (baseline, two parts of the delay, and recall). These averaged values were then analyzed using an LMEM. The LMEM showed (Fig. 4C, see Suppl. Table S4 for further details) that PLV increased for both conditions during the two parts of the delay and recall compared to the baseline (main effect of task period Delay 1 t(6032) = 12.16, p < 0.001, task period Delay 2 t(6032) = 14.13, p < 0.001, task period Recall t(6032) = 14.8, p < 0.001). The main effect of condition and the interaction between each task period and condition were not significant.

### Functional coupling between IPL and HIP

For all significant IPL-HIP connections (134 channel pairs in 6 patients), we calculated the PLV separately for each condition and first averaged the PLV over the 2-4 Hz range for each task period (baseline, two parts of the delay, and recall). The LMEM showed (Fig. 4D, see Suppl. Table S4 for further details) that PLV increased for both conditions during the two parts of the delay compared to the baseline (main effect of task period Delay 1 t(1064) = 9.17, p < 0.001, task period Delay 2 t(1064) = 8.35, p < 0.001). Additionally, there was a significant difference between the same and different objects conditions during the first half of the delay and the recall. During the first half of the delay, PLV for the different condition was higher, while during the recall, PLV for the same condition was higher (interaction between task period Delay 1 and condition t(1064) = -3.19, p = 0.002, interaction between task period Recall and condition t(1064) = 2.57, p = 0.010).

Next, we analyzed the IPL-HIP connections in the 7-8 Hz range, which also had the above-chance proportion of significant channel pairs. The LMEM for this frequency range showed (Fig. 4D) that the main effect of condition and the interaction between each task period and condition were not significant. However, for both conditions, PLV increased during the two parts of the delay compared to the baseline (main effect of task period Delay 1 t(1064) = 6.05, p < 0.001, task period Delay 2 t(1064) = 4.53, p < 0.001).

## Discussion

Our study aimed to investigate the neural dynamics and functional connectivity between the dorsal and ventral streams during memory-guided actions using iEEG. We specifically focused on how different types of visual information (object identity and location) are processed and utilized in the two streams. First, as we hypothesized, we found sustained activity in both the dorsal (IPL) and ventral (VTC) streams throughout the delay phase, reflected by increased alpha power (8-13 Hz) for both same and different objects conditions. In contrast, theta power (2-7 Hz) increased only in the ventral stream and not during delay phase: it was higher than baseline during recall in the VTC and during both encoding and recall in the hippocampus. Second, we also hypothesized that increased synchronization between IPL and VTC during delay would indicate integrated processing of spatial information. This was supported by the increased PLV between IPL and VTC in the 2-5 Hz range throughout the delay and recall periods for both conditions. Third, we predicted additional activity in the hippocampus and stronger IPL-hippocampus synchronization during the delay in the “different objects” condition, which involves remembering both object location and identity information. Our prediction was supported by an increase in alpha power in the hippocampus only for the different objects condition, and by increased PLV between IPL and hippocampus in the 2-4 Hz and 7-8 Hz ranges, with a notable difference between the same and different conditions in the 2-4 Hz range.

Our finding of increased alpha power in both the IPL and VTC throughout the delay challenges the original “dorsal amnesia hypothesis” stating that the dorsal stream lacks a memory for visual information [2]. Our results support the revised view of this hypothesis, which suggests that with additional input from the ventral stream, the dorsal stream can support memory-guided actions [7]. Previous studies with similar tasks, but using only perceptual spatial judgments without a delay phase, showed activation in the superior and inferior parietal lobules, i.e. in the dorsal stream [35–37] (see also review [22]). In contrast, in the current task using memory-guided actions, we found activation in both the dorsal and ventral streams. These areas most likely maintain spatial information, i.e., the location of the object that is closer to the central cross, as suggested by no differences between the same and different objects conditions during the delay period in IPL and VTC. The observed ventral stream activation may also be related to the conversion of transient egocentric into stable allocentric representations. Distance estimation in our task was likely performed using egocentric coordinates, as the cross was always centered on the screen and served as a stable reference point, aligning with the participant’s midsagittal plane. Both the original and revised models link the dorsal and ventral streams with egocentric and allocentric representations, respectively. Egocentric representations, associated with the dorsal stream, are transient and support real-time actions, while allocentric representations, linked to the ventral stream, are long-lasting and serve as a basis for memory-guided actions [2, 38]. Behavioral studies seem to support this hypothesis, showing that egocentric representations degrade after a few seconds of memory delay. For example, memory-guided pointing [39] and reaching [40] were performed worse in the egocentric frame than in the allocentric reference frame in healthy subjects. Over time, egocentric representations may transform into allocentric ones, activating the ventral stream. This is indirectly supported by a case study where a patient with bilateral occipito-parietal damage was impaired only in the egocentric condition with a 1.5 second delay but not with a 5 second delay [41].

The increased alpha activity during the delay and the observed increase in theta power during encoding and recall highlight the dynamic role of brain oscillations in memory processes. The observed increase in alpha amplitude aligns with other studies showing similar increases during working memory tasks [14–16, 42]. Although alpha oscillations have often been interpreted as reflecting cortical idling [43], recent studies suggest that increased alpha amplitude may be associated with active inhibition of task-irrelevant information and attentional processes [44–46]. Increased alpha power with memory load in working memory tasks further challenges the idling hypothesis and may suggest that alpha band oscillations are directly involved in memory maintenance [45, 47]. In line with these studies, the increase in alpha power in our task likely reflects either memory maintenance or active inhibition of irrelevant information, enhancing processing of relevant spatial information in the dorsal and ventral streams. On the other hand, the observed increase in theta power during encoding and recall, particularly in the hippocampus, is likely associated with memory formation and successful retrieval [48–51]. Although in our study, we analyzed only correct trials (excluding trials with incorrect joystick responses for the object position and incorrect button responses for the object identity) as the number of the incorrect trials was too small for a meaningful analysis, we suggest that theta activity supported the initial processing and integration of visual information about the objects and their spatial relations, facilitating the formation of a robust memory representation for successful retrieval during recall.

Next, as we hypothesized, we found a significant increase in PLV between IPL and VTC during the delay, but also during recall period in the slow theta (2-5 Hz) frequency range, indicating robust functional connectivity between the dorsal and ventral streams during memory-guided actions. This low-frequency synchronization suggests long-range communication necessary for the integration of spatial object information across these cortical areas. Previous research has shown that oscillations in the slow theta frequency range facilitate the coordination of distant brain regions for cognitive functions, including working memory and attention [15, 18–21]. The lack of a significant difference in PLV between the same and different objects conditions suggests that the IPL-VTC connectivity primarily supports the spatial aspect of the task, regardless of the complexity introduced by different object identities. This is consistent with the hypothesis that the dorsal and ventral streams collaboratively process spatial information for guiding delayed actions, confirming a more integrated view of the two-streams model proposed by Schenk and Hesse [7]. This integrated model is further supported by findings from a fMRI study, which emphasizes large-scale functional integration rather than strict functional dissociation between the dorsal and ventral streams [52].

Our data support the hypothesis of the additional activity in the hippocampus during the delay phase in the different objects condition, which might integrate and maintain information about the object’s identity and position. Such a role is supported by our finding of an increase in alpha power in the hippocampus only for the different objects condition. Although reaction time and accuracy of reaching responses did not differ between the same and different objects conditions, the different condition was more challenging, requiring the memory of both the position and identity of the object. This aligns with studies on working memory capacity, showing that higher task demands lead to stronger medial temporal lobe involvement, particularly in the hippocampus [15, 53, 54]. Previous single-cell recordings have revealed that hippocampal maintenance neurons increase firing rates with higher workloads in both verbal and visual working memory tasks [15, 54]. Although we did not record from single neurons, the population-level alpha power increase throughout delay aligns with findings that alpha power increases with memory load [45, 47]. Interestingly, for the different condition, the alpha power was still increased in the hippocampus and also in the IPL during recall, with a decrease in alpha power for the same condition. This decrease in alpha amplitude during retrieval is commonly observed [44, 45]. The increased alpha power for the different condition during the recall/action phase possibly reflects additional maintenance of object identity information, which was recalled during the last 2 seconds of the trial when a two-alternative question about the object’s identity appeared.

The hypothesis of stronger IPL-hippocampal synchronization during the delay in the different objects condition is supported by our findings, particularly in the slow theta range (2-4 Hz). In contrast, no difference between conditions was observed in the PLV in the fast theta range (7-8 Hz). During the first half of the delay, synchronization in the slow theta range was higher for the different condition. This difference in PLV suggests condition-dependent functional connectivity, with the hippocampus playing a crucial role in integrating both spatial and object identity information at the beginning of the delay phase. This finding aligns with other studies highlighting the hippocampus’s role in binding different types of information held in memory [13, 55]. Unexpectedly, during recall, we observed a higher PLV for the same objects condition. It can be speculated that increased communication between the IPL and hippocampus may facilitate efficient retrieval of spatial information without interference from object identity information. In the fast theta range, PLV between the IPL and hippocampus was significantly higher during the entire delay for both conditions, with no significant difference between them. The distinct roles of slow and fast theta oscillations in our task highlight the functional differentiation within the theta band. Slow theta oscillations may be more involved in integrating and processing specific task-related information, while fast theta oscillations might support broader cognitive functions required for sustained task engagement. This distinction is supported by several studies showing different characteristics of slow and fast theta [56, 57]. For instance, an iEEG study showed that slow theta in the human hippocampus had higher power during successful memory encoding and was functionally related to gamma oscillations, but this result was not observed for fast theta [56]. Also, slow theta oscillations in the human hippocampus appear to be similar to the memory-related theta oscillations observed in animals [58].

Despite its strengths, our study has several limitations. The sample size of nine epilepsy patients, while typical for iEEG studies, may limit the generalizability of the findings to a broader population. A significant limitation of iEEG studies is the variation in implantation schemes across patients. In this study, more electrodes were implanted in the right hemisphere, and different hemispheres were used in PLV analysis among patients, potentially influencing the results. However, we consider our results to be valid and reliable for several reasons. First, iEEG provides high temporal and spatial resolution, offering detailed insights into neural dynamics that are not accessible through methods like fMRI or EEG. The high signal-to-noise ratio of iEEG recordings allows statistically significant results to be obtained in individual subjects, and reproducible results from the same brain structures across a few subjects are sufficient for scientific reports [59]. In our study, each conclusion is based on data from at least six patients, providing a solid foundation for our results. Second, the consistency of our findings with established theories and previous research supports their validity despite the small sample size and patient-specific factors. Future research could include a larger sample size and consider different types of memory-guided tasks to further validate and extend our findings.

In conclusion, our results advance the understanding of visual information processing in the brain and further clarify the two streams model. The findings are consistent with recent criticisms by Schenk and Hesse [7], suggesting a more integrated function of these streams. In addition to previous fMRI studies suggesting such integration, our iEEG data provide direct electrophysiological evidence of synchronized activity between these regions during memory-guided actions, emphasizing the collaborative processing of spatial and object identity information. These findings on the interaction between dorsal and ventral streams may have potential clinical implications, particularly for patients with visuomotor impairments. For instance, neurofeedback and brain stimulation techniques could be developed to enhance synchronization between the dorsal and ventral streams, potentially improving visuomotor coordination and performance.

## Data availability

The data from this study will be available on request from the corresponding author. The codes used to produce the results in the paper are freely available at the repository https://github.com/kamilvlcek/iEEG_scripts/tree/v3.0.0

## Acknowledgments

The research was supported by project no. LX22NPO5107 (MEYS): Financed by European Union – Next Generation EU, the Czech Science Foundation (Grant No. 20-21339S), the Grant Agency of Charles University (GAUK Grant No. 248122 and 272221), ERDF-Project Brain dynamics, No. CZ.02.01.01/00/22_008/0004643, and the Ministry of Health of the Czech Republic project NU21J-08-00081. The authors are grateful to CESNET for access to their data storage facility. We would also like to thank all the patients who participated in this study.

## References

[1] Goodale MA, Milner AD. Separate visual pathways for perception and action. Trends Neurosci 1992, 15: 20–25.

[2] Goodale MA, Westwood DA, Milner AD. Two distinct modes of control for object-directed action. The roots of visual awareness: a festschrift in honour of Alan Cowey 2004: 131–144.

[3] Milner AD, Goodale MA. Two visual systems re-viewed. Neuropsychologia 2008, 46: 774–785.

[4] Milner AD, Paulignan, Y., Dijkerman, H. C., Michel, F., & Jeannerod, M. A paradoxical improvement of misreaching in optic ataxia: new evidence for two separate neural systems for visual localization. Proceedings. Biological sciences 1999.

[5] Singhal A, Monaco S, Kaufman LD, Culham JC. Human fMRI reveals that delayed action re-recruits visual perception. PLoS One 2013, 8: e73629.

[6] Cohen NR, Cross ES, Tunik E, Grafton ST, Culham JC. Ventral and dorsal stream contributions to the online control of immediate and delayed grasping: a TMS approach. Neuropsychologia 2009, 47: 1553–1562.

[7] Schenk T, Hesse C. Do we have distinct systems for immediate and delayed actions? A selective review on the role of visual memory in action. Cortex 2018, 98: 228–248.

[8] Himmelbach M, Nau M, Zundorf I, Erb M, Perenin MT, Karnath HO. Brain activation during immediate and delayed reaching in optic ataxia. Neuropsychologia 2009, 47: 1508–1517.

[9] Fiehler K, Bannert MM, Bischoff M, Blecker C, Stark R, Vaitl D, et al. Working memory maintenance of grasp-target information in the human posterior parietal cortex. Neuroimage 2011, 54: 2401–2411.

[10] Hutchison RM, Gallivan JP. Functional coupling between frontoparietal and occipitotemporal pathways during action and perception. Cortex 2018, 98: 8–27.

[11] Yonelinas AP. The hippocampus supports high-resolution binding in the service of perception, working memory and long-term memory. Behav Brain Res 2013, 254: 34–44.

[12] Ekstrom AD, Yonelinas AP. Precision, binding, and the hippocampus: Precisely what are we talking about? Neuropsychologia 2020, 138: 107341.

[13] Borders AA, Ranganath C, Yonelinas AP. The hippocampus supports high-precision binding in visual working memory. Hippocampus 2022, 32: 217–230.

[14] Hsieh LT, Ekstrom AD, Ranganath C. Neural oscillations associated with item and temporal order maintenance in working memory. J Neurosci 2011, 31: 10803–10810.

[15] Boran E, Fedele T, Klaver P, Hilfiker P, Stieglitz L, Grunwald T, et al. Persistent hippocampal neural firing and hippocampal-cortical coupling predict verbal working memory load. Sci Adv 2019, 5: eaav3687.

[16] Lee B, Kim JS, Chung CK. Parietal and medial temporal lobe interactions in working memory goal-directed behavior. Cortex 2022, 150: 126–136.

[17] Su M, Hu K, Liu W, Wu Y, Wang T, Cao C, et al. Theta Oscillations Support Prefrontal-hippocampal Interactions in Sequential Working Memory. Neurosci Bull 2024, 40: 147–156.

[18] Fries P. Rhythms for Cognition: Communication through Coherence. Neuron 2015, 88: 220–235.

[19] Pesaran B, Vinck M, Einevoll GT, Sirota A, Fries P, Siegel M, et al. Investigating large-scale brain dynamics using field potential recordings: analysis and interpretation. Nat Neurosci 2018, 21: 903–919.

[20] Sarnthein J, Petsche, H., Rappelsberger, P., Shaw, G. L., & von Stein, A. Synchronization between prefrontal and posterior association cortex during human working memory. Proceedings of the National Academy of Sciences of the United States of America 1998, 95: 7092–7096.

[21] Solomon EA, Kragel JE, Sperling MR, Sharan A, Worrell G, Kucewicz M, et al. Widespread theta synchrony and high-frequency desynchronization underlies enhanced cognition. Nat Commun 2017, 8: 1704.

[22] Moraresku S, Vlcek K. The use of egocentric and allocentric reference frames in static and dynamic conditions in humans. Physiol Res 2020, 69: 787–801.

[23] Peirce J, Gray JR, Simpson S, MacAskill M, Hochenberger R, Sogo H, et al. PsychoPy2: Experiments in behavior made easy. Behav Res Methods 2019, 51: 195–203.

[24] Janca R, Tomasek M, Kalina A, Marusic P, Krsek P, Lesko R. Automated identification of stereoelectroencephalography contacts and measurement of factors influencing accuracy of frame stereotaxy. IEEE Journal of Biomedical and Health Informatics 2023, 27: 3326–3336.

[25] Vlcek K, Fajnerova I, Nekovarova T, Hejtmanek L, Janca R, Jezdik P, et al. Mapping the Scene and Object Processing Networks by Intracranial EEG. Front Hum Neurosci 2020, 14: 561399.

[26] Moraresku S, Hammer J, Janca R, Jezdik P, Kalina A, Marusic P, et al. Timing of Allocentric and Egocentric Spatial Processing in Human Intracranial EEG. Brain Topogr 2023, 36: 870–889.

[27] Gunia A, Moraresku S, Janca R, Jezdik P, Kalina A, Hammer J, et al. The brain dynamics of visuospatial perspective-taking captured by intracranial EEG. Neuroimage 2024, 285: 120487.

[28] Miller KJ, Honey CJ, Hermes D, Rao RP, denNijs M, Ojemann JG. Broadband changes in the cortical surface potential track activation of functionally diverse neuronal populations. Neuroimage 2014, 85 Pt 2: 711–720.

[29] Janca R, Jezdik P, Cmejla R, Tomasek M, Worrell GA, Stead M, et al. Detection of interictal epileptiform discharges using signal envelope distribution modelling: application to epileptic and non-epileptic intracranial recordings. Brain Topogr 2015, 28: 172–183.

[30] Lachaux JP, Rodriguez E, Martinerie J, Varela FJ. Measuring phase synchrony in brain signals. Hum Brain Mapp 1999, 8: 194–208.

[31] Oostenveld R, Fries P, Maris E, Schoffelen JM. FieldTrip: Open source software for advanced analysis of MEG, EEG, and invasive electrophysiological data. Comput Intell Neurosci 2011, 2011: 156869.

[32] Dimakopoulos V, Megevand P, Stieglitz LH, Imbach L, Sarnthein J. Information flows from hippocampus to auditory cortex during replay of verbal working memory items. Elife 2022, 11.

[33] Park YM, Park J, Baek JH, Kim SI, Kim IY, Kang JK, et al. Differences in theta coherence between spatial and nonspatial attention using intracranial electroencephalographic signals in humans. Hum Brain Mapp 2019, 40: 2336–2346.

[34] Dempsey LA, Cooper RJ, Roque T, Correia T, Magee E, Powell S, et al. Data-driven approach to optimum wavelength selection for diffuse optical imaging. J Biomed Opt 2015, 20: 016003.

[35] Galati G, Lobel E, Vallar G, Berthoz A, Pizzamiglio L, Le Bihan D. The neural basis of egocentric and allocentric coding of space in humans: a functional magnetic resonance study. Exp Brain Res 2000, 133: 156–164.

[36] Saj A, Cojan Y, Musel B, Honore J, Borel L, Vuilleumier P. Functional neuro-anatomy of egocentric versus allocentric space representation. Neurophysiol Clin 2014, 44: 33–40.

[37] Ruotolo F, Ruggiero G, Raemaekers M, Iachini T, van der Ham IJM, Fracasso A, et al. Neural correlates of egocentric and allocentric frames of reference combined with metric and non-metric spatial relations. Neuroscience 2019, 409: 235–252.

[38] Goodale MA, Haffenden A. Frames of reference for perception and action in the human visual system. Neuroscience & Biobehavioral Reviews 1998, 22: 161–172.

[39] Hay L, Redon C. Response delay and spatial representation in pointing movements. Neurosci Lett 2006, 408: 194–198.

[40] Chen Y, Byrne P, Crawford JD. Time course of allocentric decay, egocentric decay, and allocentric-to-egocentric conversion in memory-guided reach. Neuropsychologia 2011, 49: 49–60.

[41] Ilardi CR, Iavarone A, Villano I, Rapuano M, Ruggiero G, Iachini T, et al. Egocentric and allocentric spatial representations in a patient with Balint-like syndrome: A single-case study. Cortex 2021, 135: 10–16.

[42] Johnson EL, King-Stephens D, Weber PB, Laxer KD, Lin JJ, Knight RT. Spectral Imprints of Working Memory for Everyday Associations in the Frontoparietal Network. Front Syst Neurosci 2018, 12: 65.

[43] Pfurtscheller G, Stancak Jr A, Neuper C. Event-related synchronization (ERS) in the alpha band—an electrophysiological correlate of cortical idling: a review. International journal of psychophysiology 1996, 24: 39–46.

[44] Sauseng P, Klimesch W, Doppelmayr M, Pecherstorfer T, Freunberger R, Hanslmayr S. EEG alpha synchronization and functional coupling during top-down processing in a working memory task. Hum Brain Mapp 2005, 26: 148–155.

[45] Klimesch W, Sauseng P, Hanslmayr S. EEG alpha oscillations: the inhibition-timing hypothesis. Brain Res Rev 2007, 53: 63–88.

[46] Klimesch W. alpha-band oscillations, attention, and controlled access to stored information. Trends Cogn Sci 2012, 16: 606–617.

[47] Jensen O, Gelfand J, Kounios J, Lisman JE. Oscillations in the alpha band (9–12 Hz) increase with memory load during retention in a short-term memory task. Cerebral cortex 2002, 12: 877–882.

[48] Sederberg PB, Kahana MJ, Howard MW, Donner EJ, Madsen JR. Theta and gamma oscillations during encoding predict subsequent recall. Journal of Neuroscience 2003, 23: 10809–10814.

[49] Caplan JB, Glaholt MG. The roles of EEG oscillations in learning relational information. Neuroimage 2007, 38: 604–616.

[50] Herweg NA, Solomon EA, Kahana MJ. Theta Oscillations in Human Memory. Trends Cogn Sci 2020, 24: 208–227.

[51] Joensen BH, Bush D, Vivekananda U, Horner AJ, Bisby JA, Diehl B, et al. Hippocampal theta activity during encoding promotes subsequent associative memory in humans. Cereb Cortex 2023, 33: 8792–8802.

[52] Ray D, Hajare N, Roy D, Banerjee A. Large-scale Functional Integration, Rather than Functional Dissociation along Dorsal and Ventral Streams, Underlies Visual Perception and Action. J Cogn Neurosci 2020, 32: 847–861.

[53] Axmacher N, Mormann F, Fernandez G, Cohen MX, Elger CE, Fell J. Sustained neural activity patterns during working memory in the human medial temporal lobe. J Neurosci 2007, 27: 7807–7816.

[54] Boran E, Hilfiker P, Stieglitz L, Sarnthein J, Klaver P. Persistent neuronal firing in the medial temporal lobe supports performance and workload of visual working memory in humans. Neuroimage 2022, 254: 119123.

[55] Olson IR, Page K, Moore KS, Chatterjee A, Verfaellie M. Working memory for conjunctions relies on the medial temporal lobe. J Neurosci 2006, 26: 4596–4601.

[56] Lega BC, Jacobs J, Kahana M. Human hippocampal theta oscillations and the formation of episodic memories. Hippocampus 2012, 22: 748–761.

[57] Lisman JE, Jensen O. The theta-gamma neural code. Neuron 2013, 77: 1002–1016.

[58] Watrous AJ, Lee DJ, Izadi A, Gurkoff GG, Shahlaie K, Ekstrom AD. A comparative study of human and rat hippocampal low-frequency oscillations during spatial navigation. Hippocampus 2013, 23: 656–661.

[59] Lachaux JP, Rudrauf D, Kahane P. Intracranial EEG and human brain mapping. J Physiol Paris 2003, 97: 613–628.

