## Supplementary Material for "Neural dynamics of visual streams interactions during memory-guided actions investigated by intracranial EEG"

### Table S1. Subject information.

| Patient | Age | Gender | Handedness | Education | Epilepsy duration (years) | Suspected seizure zone | Epilepsy pathology |
|---|---|---|---|---|---|---|---|
| P1 | 43 | M | R | primary | 33 | R temporal and parietal lobes | suspected FCD I |
| P2 | 37 | F | L | tertiary | 13 | bilateral temporal lobes | FCD I |
| P3 | 36 | M | R | tertiary | 12 | R temporal and parietal lobes, insula | post-infectious gliosis |
| P4 | 24 | F | L | primary | 8 | R temporal lobe | periventricular nodular heterotopy |
| P5 | 49 | M | R | tertiary | 27 | bilateral temporal and parietal lobes | hippocampal sclerosis |
| P6 | 30 | F | L | tertiary | 19 | L temporal lobe | periventricular nodular heterotopy |
| P7 | 46 | M | R | primary | 36 | R occipital and parietal lobes | suspected FCD I |
| P8 | 30 | M | L | primary | 28 | R operculo-insular and temporal lobe | FCD I |
| P9 | 36 | F | R | tertiary | 7 | R temporal, parietal, and occipital lobes | suspected FCD II |

F, female; M, male; R, right; L, left; FCD, Focal Cortical Dysplasia

**Table S2. Linear mixed effects model results for alpha power (8-13 Hz).**

**IPL**

| fixed effect | Estimate | SE | DF | tStat | pValue |
|---|---|---|---|---|---|
| task period Encoding | -0.034 | 0.033 | 430 | -1.03 | 0.3028 |
| task period Delay 1 | 0.137 | 0.033 | 430 | 4.16 | 0.0000 * |
| task period Delay 2 | 0.171 | 0.033 | 430 | 5.18 | 0.0000 * |
| task period Recall | 0.042 | 0.033 | 430 | 1.28 | 0.2012 |
| condition Same | 0.000 | 0.033 | 430 | -0.01 | 0.9897 |
| task period Encoding : condition Same | 0.001 | 0.047 | 430 | 0.03 | 0.9750 |
| task period Delay 1 : condition Same | -0.071 | 0.047 | 430 | -1.52 | 0.1305 |
| task period Delay 2 : condition Same | -0.116 | 0.047 | 430 | -2.47 | 0.0137 |
| task period Recall : condition Same | -0.147 | 0.047 | 430 | -3.14 | 0.0018 * |

**VTC**

| fixed effect | Estimate | SE | DF | tStat | pValue |
|---|---|---|---|---|---|
| task period Encoding | -0.130 | 0.025 | 630 | -5.23 | 0.0000 * |
| task period Delay 1 | 0.111 | 0.025 | 630 | 4.46 | 0.0000 * |
| task period Delay 2 | 0.160 | 0.025 | 630 | 6.48 | 0.0000 * |
| task period Recall | -0.024 | 0.025 | 630 | -0.97 | 0.3313 |
| condition Same | 0.000 | 0.025 | 630 | 0.01 | 0.9937 |
| task period Encoding : condition Same | 0.031 | 0.035 | 630 | 0.89 | 0.3726 |
| task period Delay 1 : condition Same | 0.000 | 0.035 | 630 | 0.01 | 0.9946 |
| task period Delay 2 : condition Same | -0.030 | 0.035 | 630 | -0.85 | 0.3930 |
| task period Recall : condition Same | -0.060 | 0.035 | 630 | -1.70 | 0.0889 |

**HIP**

| fixed effect | Estimate | SE | DF | tStat | pValue |
|---|---|---|---|---|---|
| task period Encoding | 0.056 | 0.032 | 190 | 1.72 | 0.0868 |
| task period Delay 1 | 0.117 | 0.032 | 190 | 3.63 | 0.0004 * |
| task period Delay 2 | 0.071 | 0.032 | 190 | 2.19 | 0.0295 |
| task period Recall | 0.105 | 0.032 | 190 | 3.24 | 0.0014 * |
| condition Same | -0.001 | 0.032 | 190 | -0.03 | 0.9723 |
| task period Encoding : condition Same | -0.117 | 0.046 | 190 | -2.57 | 0.0110 |
| task period Delay 1 : condition Same | -0.209 | 0.046 | 190 | -4.57 | 0.0000 * |
| task period Delay 2 : condition Same | -0.170 | 0.046 | 190 | -3.73 | 0.0003 * |
| task period Recall : condition Same | -0.160 | 0.046 | 190 | -3.51 | 0.0006 * |

* significant after Bonferroni correction, $p < 0.008$; SE, standard error; DF, degrees of freedom

**Table S3. Linear mixed effects model results for theta power (2-7 Hz).**

**IPL**

| fixed effect | Estimate | SE | DF | tStat | pValue |
|---|---|---|---|---|---|
| task period Encoding | 0.078 | 0.045 | 300 | 1.70 | 0.0894 |
| task period Delay 1 | 0.058 | 0.045 | 300 | 1.27 | 0.2064 |
| task period Delay 2 | 0.052 | 0.045 | 300 | 1.13 | 0.2581 |
| task period Recall | 0.096 | 0.045 | 300 | 2.12 | 0.0350 |
| condition Same | 0.000 | 0.045 | 300 | 0.01 | 0.9929 |
| task period Encoding : condition Same | -0.025 | 0.064 | 300 | -0.39 | 0.6979 |
| task period Delay 1 : condition Same | -0.069 | 0.064 | 300 | -1.07 | 0.2847 |
| task period Delay 2 : condition Same | -0.016 | 0.064 | 300 | -0.25 | 0.8042 |
| task period Recall : condition Same | -0.070 | 0.064 | 300 | -1.08 | 0.2790 |

**VTC**

| fixed effect | Estimate | SE | DF | tStat | pValue |
|---|---|---|---|---|---|
| task period Encoding | 0.076 | 0.045 | 280 | 1.69 | 0.0920 |
| task period Delay 1 | 0.099 | 0.045 | 280 | 2.20 | 0.0283 |
| task period Delay 2 | 0.106 | 0.045 | 280 | 2.35 | 0.0197 |
| task period Recall | 0.158 | 0.045 | 280 | 3.51 | 0.0005 * |
| condition Same | 0.001 | 0.045 | 280 | 0.01 | 0.9884 |
| task period Encoding : condition Same | -0.104 | 0.064 | 280 | -1.63 | 0.1034 |
| task period Delay 1 : condition Same | -0.162 | 0.064 | 280 | -2.54 | 0.0116 |
| task period Delay 2 : condition Same | -0.147 | 0.064 | 280 | -2.31 | 0.0218 |
| task period Recall : condition Same | -0.100 | 0.064 | 280 | -1.57 | 0.1166 |

**HIP**

| fixed effect | Estimate | SE | DF | tStat | pValue |
|---|---|---|---|---|---|
| task period Encoding | 0.177 | 0.031 | 170 | 5.63 | 0.0000 * |
| task period Delay 1 | 0.034 | 0.031 | 170 | 1.08 | 0.2825 |
| task period Delay 2 | 0.015 | 0.031 | 170 | 0.49 | 0.6269 |
| task period Recall | 0.154 | 0.031 | 170 | 4.90 | 0.0000 * |
| condition Same | -0.002 | 0.031 | 170 | -0.05 | 0.9563 |
| task period Encoding : condition Same | -0.043 | 0.045 | 170 | -0.97 | 0.3328 |
| task period Delay 1 : condition Same | 0.052 | 0.045 | 170 | 1.17 | 0.2437 |
| task period Delay 2 : condition Same | 0.026 | 0.045 | 170 | 0.58 | 0.5611 |
| task period Recall : condition Same | 0.053 | 0.045 | 170 | 1.19 | 0.2346 |

* significant after Bonferroni correction, $p < 0.008$; SE, standard error; DF, degrees of freedom

**Table S4. Linear mixed effects model results for PLV.**

**IPL-VTC, 2-5Hz**

| fixed effect | Estimate | SE | DF | tStat | pValue |
|---|---|---|---|---|---|
| task period Delay 1 | 0.034 | 0.003 | 6032 | 12.16 | 0.0000 * |
| task period Delay 2 | 0.040 | 0.003 | 6032 | 14.13 | 0.0000 * |
| task period Recall | 0.041 | 0.003 | 6032 | 14.79 | 0.0000 * |
| condition Same | -0.006 | 0.003 | 6032 | -2.14 | 0.0327 |
| task period Delay 1 : condition Same | 0.000 | 0.004 | 6032 | -0.12 | 0.9076 |
| task period Delay 2 : condition Same | 0.000 | 0.004 | 6032 | -0.07 | 0.9479 |
| task period Recall : condition Same | -0.002 | 0.004 | 6032 | -0.60 | 0.5510 |

**IPL-HIP, 2-4Hz**

| fixed effect | Estimate | SE | DF | tStat | pValue |
|---|---|---|---|---|---|
| task period Delay 1 | 0.064 | 0.007 | 1064 | 9.17 | 0.0000 * |
| task period Delay 2 | 0.058 | 0.007 | 1064 | 8.35 | 0.0000 * |
| task period Recall | 0.025 | 0.007 | 1064 | 3.57 | 0.0004 * |
| condition Same | 0.002 | 0.007 | 1064 | 0.36 | 0.7202 |
| task period Delay 1 : condition Same | -0.031 | 0.010 | 1064 | -3.19 | 0.0015 * |
| task period Delay 2 : condition Same | -0.014 | 0.010 | 1064 | -1.44 | 0.1503 |
| task period Recall : condition Same | 0.025 | 0.010 | 1064 | 2.57 | 0.0104 * |

**IPL-HIP, 7-8Hz**

| fixed effect | Estimate | SE | DF | tStat | pValue |
|---|---|---|---|---|---|
| task period Delay 1 | 0.043 | 0.007 | 1064 | 6.05 | 0.0000 * |
| task period Delay 2 | 0.032 | 0.007 | 1064 | 4.53 | 0.0000 * |
| task period Recall | 0.009 | 0.007 | 1064 | 1.22 | 0.2215 |
| condition Same | 0.004 | 0.007 | 1064 | 0.61 | 0.5444 |
| task period Delay 1 : condition Same | -0.009 | 0.010 | 1064 | -0.86 | 0.3894 |
| task period Delay 2 : condition Same | -0.001 | 0.010 | 1064 | -0.09 | 0.9252 |
| task period Recall : condition Same | -0.012 | 0.010 | 1064 | -1.16 | 0.2476 |

* significant after Bonferroni correction, $p < 0.0167$; SE, standard error; DF, degrees of freedom
